# Kinetic mechanism of human mitochondrial RNase P

**DOI:** 10.1101/666792

**Authors:** Xin Liu, Nancy Wu, Aranganathan Shanmuganathan, Bradley P. Klemm, Michael J. Howard, Wan Hsin Lim, Markos Koutmos, Carol A. Fierke

## Abstract

A first step in processing mitochondrial precursor tRNA (pre-tRNA) is cleavage of the 5’ leader catalyzed by ribonuclease P (RNase P). Human mitochondrial RNase P (mtRNase P) is composed of three protein subunits: mitochondrial RNase P protein (MRPP) 1, 2 and 3. Even though MRPP3 is the metallonuclease subunit responsible for catalysis, cleavage is observed only in the presence of the MRPP1/2 subcomplex. To understand the functional role of MRPP1/2, we reconstituted human mitochondrial RNase P *in vitro* and performed kinetic and thermodynamic analyses. MRPP1/2 significantly enhances both the catalytic activity and the apparent substrate affinity of mtRNase P. Additionally, pull-down and binding data demonstrate synergy between binding pre-tRNA and formation of a catalytically active MRPP1/2/3 complex. These data suggest that conformational changes in the MRPP1/2-pre-tRNA complex lead to protein-protein or protein-RNA interactions that increase both MRPP3 recognition and cleavage efficiency. This work presents the first kinetic model for human mtRNase P, providing a fundamental framework for the function of MRPP1/2 for recognition and processing of pre-tRNA.

---

Dysfunction in mitochondria, the powerhouses of cells, is increasingly implicated in human aging, neurodegenerative diseases, cancer, diabetes, and various rare diseases (1). While the etiology of many of these diseases remains unknown, mitochondrial dysfunction is frequently linked to damage and mutations in the mitochondrial genome. About half of these disease-related point mutations map to mitochondrial tRNA (mt-tRNA) genes (2-5). The human mitochondrial genome is transcribed into three polycistronic transcripts, which are first processed by nuclear-encoded mitochondrial RNase P and RNase Z to generate tRNA with matured 5’ and 3’ ends, respectively. The combined cleavage activity of RNase P and RNase Z on the polycistronic transcripts also releases the interspersed mRNA, ribosomal RNA and long noncoding RNA transcripts that flank the excised mt-tRNA (6,7). Therefore, endonuclease processing of the mitochondrial transcript is a critical step in mitochondrial biogenesis (5,6). Understanding the mt-tRNA processing step catalyzed by human mtRNase P can provide valuable insights into important pathways and protein function in the mitochondria.

RNase P is a divalent metal ion dependent endonuclease found in all domains of life and is the only conserved step in the tRNA maturation pathway. RNase P requires magnesium ions to stabilize the structure of pre-tRNA substrates and to catalyze cleavage of the 5’ leader from pre-tRNA *in vivo* (8). In many organisms, RNase P is a ribonucleoprotein complex with a catalytic RNA subunit and accessory proteins (9,10). However, in plant mitochondria, nuclei, and chloroplasts, a protein-only RNase P (PRORP) functions as a single-subunit endonuclease (11,12). A homolog of PRORP, MRPP3, in human mitochondria functions in a multi-subunit complex with MRPP1 and MRPP2 (13). MRPP3 is the catalytic subunit containing the active site for catalyzing pre-tRNA hydrolysis, but the endonuclease activity is dependent on the presence of MRPP1 and MRPP2 (13). The mechanistic role of each MRPP subunit for substrate recognition and catalysis is poorly understood.

MRPP1, 2, and 3 are required for the proper processing of mitochondrial transcripts *in vivo*, including transcripts that encode for respiratory chain complexes. Disruption of these processes leads to mitochondrial dysfunction and disease (14-17). Recessive MRPP1 mutations destabilize the mutant protein and lead to inadequate processing of mt-tRNA and low levels of assembled respiratory chain complexes (15). Pathogenic mutations located in MRPP2 lead to a reduction in MRPP1 and MRPP2 protein levels and tRNA processing (18,19). Both MRPP1 and MRPP3 partial knockdown experiments result in enrichment of precursor transcripts indicating a reduction in tRNA and RNA processing (6). The loss of any of the MRPP3, MRPP1 or MRPP2 in Drosophila causes aberrant tRNA processing and lethality (16) while knockout of MRPP3 is embryonically lethal in mice (17).

Apart from their roles in human mtRNase P, MRPP1, a tRNA methyltransferase, and MRPP2, a hydroxysteroid (17-β) dehydrogenase, function together in a stable complex (MRPP1/2) to catalyze *N*1-methylation of A or G at position 9 in mt-tRNA (20). MRPP2 is an NAD-dependent member of the short-chain dehydrogenase/reductase superfamily involved in the oxidation of isoleucine, steroids, and fatty acids (21,22). MRPP1 requires MRPP2 for stability and function, but the dehydrogenase activity and NAD binding capability of MRPP2 are dispensable for methyltransferase activity (13,23). MRPP2 mutations and knockdown experiments indicate that MRPP2 is required for stable expression of MRPP1 (19). Furthermore, the enzymatic activities of MRPP1 and MRPP2 have been reported to have little effect on MRPP3 cleavage activity (23). The MRPP1/2 complex was also shown to enhance the 3’ end processing by the RNase Z enzyme ELAC2 for 17 of the 22 encoded mt-tRNAs (24). Two crystal structures of MRPP3 with significant N-terminal truncations reveal a distorted active site with no bound metal ions leading to the hypothesis that MRPP1/2 may mediate metal binding at the active site by stabilizing an active conformation of MRPP3 (25,26). Alternatively, the N-terminal truncations in MRPP3 may cause the low activity, as suggested by the inability of MRPP1/2 to activate the truncated MRPP3 constructs. Thus, MRPP1 and MRPP2 are proposed to play a scaffolding role in mtRNase P that is independent of their primary catalytic functions (26). However, this proposal remains unverified by biochemical investigations into the role that MRPP1/2 plays in the catalytic pathway of mtRNase P.

Here, we report the first kinetic mechanism of human mtRNase P and uncover the role of the MRPP1/2 subcomplex using a bacterial model pre-tRNA substrate. We performed steady-state and transient kinetics, fluorescence anisotropy binding assays, and pull-down experiments to probe the role of the MRPP1/2 subcomplex. Our data demonstrate that MRPP1/2 enhances the catalytic activity of MRPP3 by increasing substrate affinity and cleavage activity. Furthermore, we find that the MRPP1/2/3 ternary complex is stabilized by pre-tRNA. Kinetic data show that a high concentration of MRPP1/2 is required to activate MRPP3 to efficiently cleave pre-tRNA, consistent with results showing that a partial MRPP1 knockdown had a larger effect on RNA transcript levels in cells compared to a similar level of MRPP3 knockdown (6). Our data support a kinetic model where MRPP1/2 undertakes a conformational selection step upon binding to pre-tRNA. This work provides a mechanistic framework to understand mt-tRNA-related diseases and will guide future work in dissecting the function of this essential multi-subunit RNase P.

## RESULTS

### MRPP1/2 enhances pre-tRNA cleavage catalyzed by MRPP3

To assess the contribution of MRPP1/2 on mtRNase P function, we performed both multiple-turnover (MTO) and single-turnover (STO) kinetics experiments. STO measurements provide insight on the kinetic pathway up to the cleavage step, while MTO kinetics extends to include every step including the product dissociation step. In our MTO experiments, we used a previously described *B. subtilis* fluorescein-labelled pre-tRNA^Asp^ (Fl-pre-tRNA) substrate with a canonical cloverleaf secondary structure and predicted tertiary fold. This Fl-pre-tRNA was used to elucidate the catalytic mechanism of a homolog of MRPP3 from *A. thaliana*, PRORP1 (27,28). We chose this substrate so we can compare the kinetics of MRPP1/2/3 with PRORP1. STO kinetics and MTO activity with only a single time-point have been previously described for human mtRNase P (13,14,23,24,29). Here, we present the first detailed MTO kinetic analysis.

MRPP1/2/3 catalyzes complete cleavage of Fl-pre-tRNA (Figure 1A) and the initial rate for cleavage depends on both pre-tRNA and MRPP1/2 concentrations (Figures 1B, C). The apparent *K*_M_ value (*K*_M,app_) for pre-tRNA is reduced by at least 4-fold, and the apparent turnover rate (*k*_cat, app_) is increased by at least 3-fold when MRPP1/2 concentrations are increased from 150 nM to 1200 nM (Table 1). At saturating concentrations of MRPP1/2 the catalytic efficiency of MRPP1/2/3 ((*k*_cat_*/K*_M_)_,app_ = 4 × 10^5^ M^−1^s^−1^, Figure 1F and Table 1) is comparable to that of *A. thaliana* PRORP1 (1 × 10^5^ M^−1^s^−1^) (27). In the absence of MRPP1/2, no significant product formation is observed under MTO or STO conditions, even after 20 hours incubation of 10 nM pre-tRNA with 25 μM MRPP3 (data not shown). Based on these data, the value of *k*_cat_*/K*_M_ catalyzed by MRPP3 alone is negligible (estimated to be less than 4 M^−1^s^−1^, Table 1).^1^ As a result, the MRPP1/2 subcomplex enhances catalytic efficiency by more than 100,000-fold (Table 1). The *K*_1/2_ for activation of MRPP3-catalyzed turnover by MRPP1/2 is 0.4 µM (Figure 1D). These results demonstrate that MRPP3 efficiently catalyzes pre-tRNA cleavage in the presence of MRPP1/2 and high concentrations of MRPP1/2 are needed for maximal MTO activity.

**Table 1.** Apparent steady-state kinetic parameters (MTO conditions) of mtRNase P activity as a function of MRPP1/2 concentration as described in Figure 1.

| MRPP1/2<br>(nM) | $k_{cat,app}$<br>(s<sup>-1</sup>) | $K_{M,app}$<br>(μM) | $(k_{cat}/K_M)_{app}$<br>(M<sup>-1</sup>s<sup>-1</sup>) |
| --- | --- | --- | --- |
| 1200 | 0.19 ± 0.01 | 0.47 ± 0.08 | (4.0 ± 0.1) × 10<sup>5</sup> |
| 500 | 0.15 ± 0.01 | 1.2 ± 0.3 | (1.2 ± 0.2) × 10<sup>5</sup> |
| 350 | 0.14 ± 0.02 | 1.7 ± 0.7 | (8 ± 2) × 10<sup>4</sup> |
| 150 | 0.059 ± 0.004 | 2.1 ± 0.3 | (2.9 ± 0.3) × 10<sup>4</sup> |
| 0 | ND | ND | < 4 |

ND: not determined.
Reactions contained MRPP3 (25–50 nM), varying concentrations of MRPP1/2 (150–1200 nM) and Fl-pre-tRNA substrate (150–5000 nM of total pre-tRNA with 40 nM of Fl-pre-tRNA and varying unlabeled pre-tRNA). Reactions were initiated by mixing Fl-pre-tRNA with MRPP proteins in 50 mM MOPS pH 7.8, 100 mM NaCl, 4.5 mM MgCl<sub>2</sub> and 1 mM TCEP at 37 °C.

**Figure 1.**
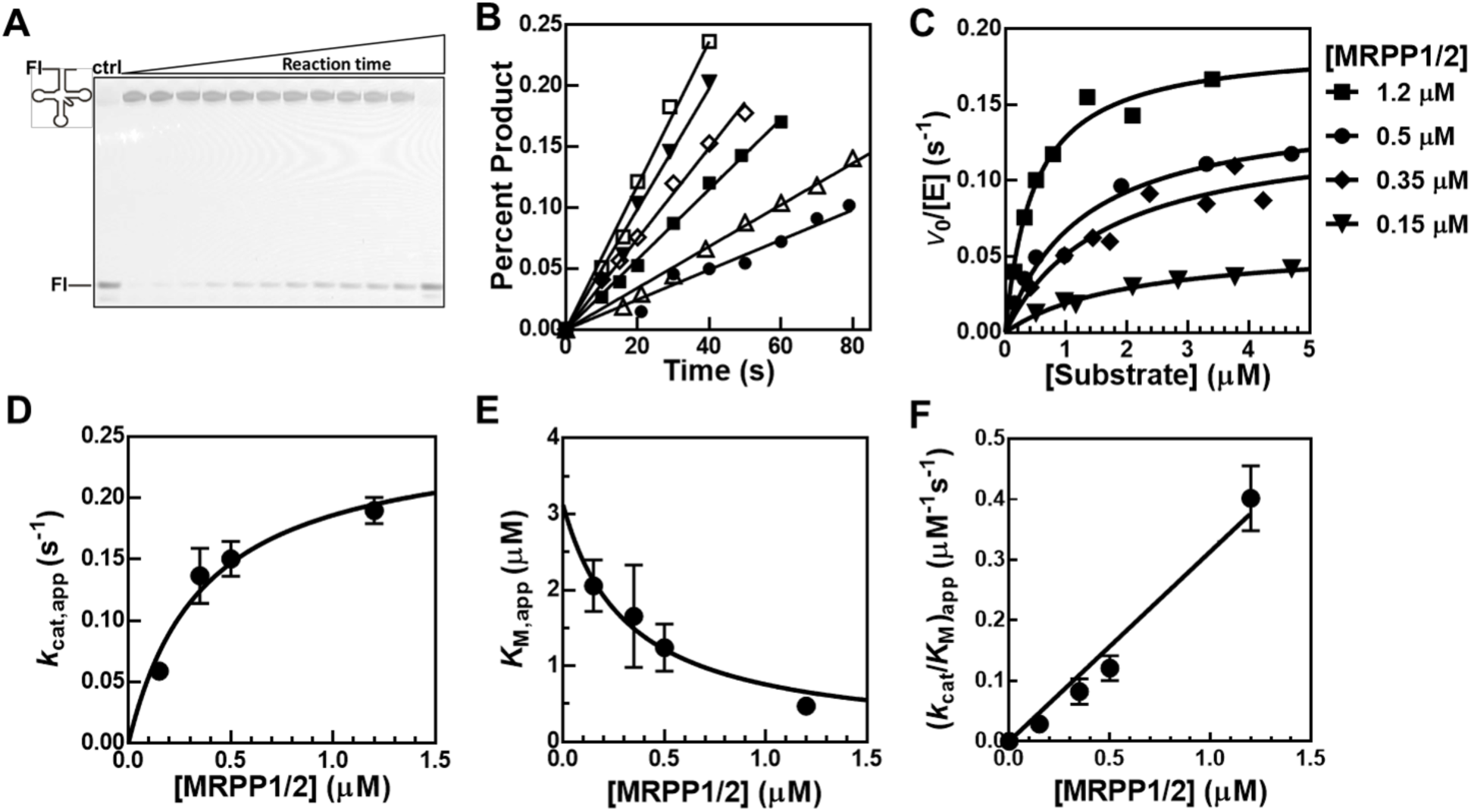
Multiple-turnover (MTO) activity of human mtRNase P. (**A**) A representative fluorescence scan (λ_ex_ = 488 nm, λ_em_ = 525 nm) of a 22% polyacrylamide gel with samples from a MTO assay containing 1200 nM MRPP1/2, 25 nM MRPP3, 1300 nM pre-tRNA (containing 40 nM Fl-pre-tRNA). Time points are: 10 s, 15 s, 20 s, 30 s, 40 s, 49 s, 1min, 1 min 11 s, 1 min 20 s, 1 min 29 s, 2 min, and 1h. Control lane is reaction of 20 nM PRORP1 with 100 nM substrate (containing 40 nM Fl-pre-tRNA) at 1h. All the reactions were performed in 50 mM MOPS pH 7.8, 100 mM NaCl, 4.5 mM MgCl_2_ and 1 mM TCEP at 37 °C. (**B**) Representative initial rates of percent cleavage under MTO conditions as described in (A) at varying substrate concentration (300 nM open square; 500 nM filled triangle; 800 nM open diamond; 1300 nM filled square; 2100 nM open triangle; 3400 nM filled circle). (**C**) The dependence of the initial velocities of MRPP activity on pre-tRNA concentration (150–4700 nM) and varying MRPP1/2 concentrations. The Michaelis-Menten equation was fit to the data to obtain steady-state kinetic parameters *k*_cat,app_, *K*_M,app_, and (*k*_cat_/*K*-_M_)_app_. The apparent kinetic parameters are listed in Table 1. (**D to F**) Dependence of kinetic parameters on MRPP1/2 concentration. Equations 1, 2 and 3 were fit independently to the data to obtain the dependence of *k*_cat,app_ (**D**), *K*_M,app_ (**E**), and (*k*_cat/_*K*_M_)_app_ (**F**) on MRPP1/2 concentration, respectively. The errors represent the standard error from fitting initial rate data of seven different substrate concentrations in (C). (**D**) and (**F**) are used to determine thermodynamic and kinetic rate constants as described in methods: *k*_chem_ = 0.25 ± 0.04 s^−1^, *K*_4_ = 0.4 ± 0.1 µM, *K*_2_*K*_5_ = 0.8 ± 0.1 µM^2^.

To determine if catalysis is the rate-limiting step in MRPP1/2/3 cleavage activity, we measured single turnover (STO) kinetics (Figure 2). The time-dependence of the STO cleavage was fit to a single exponential decay (Figure 2A). The observed rate constant (*k*_obs_) for cleavage catalyzed by MRPP3 has a hyperbolic dependence on the concentration of MRPP1/2 (Figure 2B). Additionally, increasing the MRPP3 concentration both enhances the cleavage rate constant at saturating MRPP1/2 (*k*_max,app_) and decreases the *K*_1/2_ for MRPP1/2 (*K*_1/2_^MRPP1/2^) more than 3-fold (from 1.0 to 0.3 µM) (Figure 2B), consistent with the formation of a MRPP1/2/3•pre-tRNA complex. The *K*_1/2_ for MRPP3 at saturating MRPP1/2 (*K*_1/2_^MRPP3^) is 0.094 ± 0.003 µM (Figure 2C). The STO cleavage rate constant at saturating MRPP1/2 and MRPP3 (*k*_max_) is 0.21 ± 0.01 s^−1^ (Figure 2C, Figure S3), similar to the maximum *k*_cat,app_ value (*k*_chem_) calculated from the MTO kinetics (Figure 1D, Equation 1), indicating that under MTO conditions the cleavage step (*k*_chem_) is likely rate-limiting.

**Figure 2.**
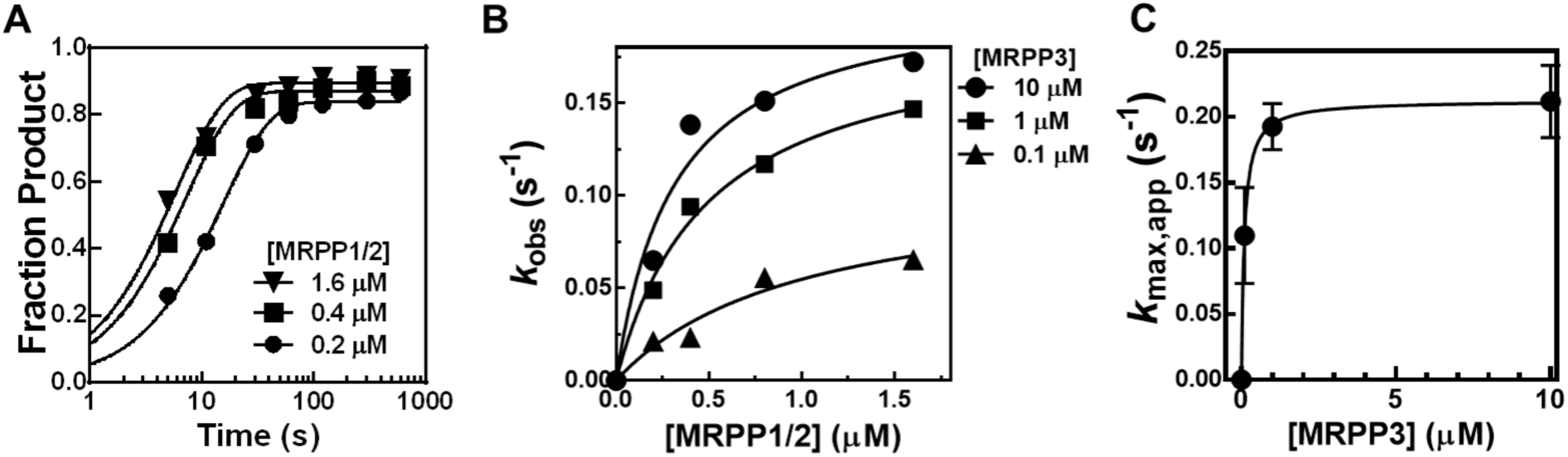
Single-turnover (STO) activity of mtRNase P. (**A**) A representative time-course of STO activity of MRPP1/2/3 complex using 10 µM MRPP3 and varying concentrations of MRPP1/2. A single exponential equation (Equation 7) was fit to the data to obtain the *k*_obs_. (**B**) A hyperbola (Equation 8) was fit to the MRPP1/2 dependence of *k*_obs_ at various concentrations of MRPP3 to determine values for *k*_max,app_ and *K*_1/2_^MRPP1/2^. (**C**) A hyperbola (Equation 8) was fit to the MRPP3 dependence of *k*_max,app_ to determine values for the apparent *K*_1/2_^MRPP3^ (0.094 ± 0.003 µM) and *k*_max_ (0.21 ± 0.001 s^−1^). The *K*_1/2_^MRPP3^ is approximately the value of *K*_6_ by rapid-equilibrium approximation under STO conditions. The reported errors and error bars are the standard error of the mean values obtained from fitting the data of five different MRPP1/2 concentrations in (B).

### Pre-tRNA substrate enhances formation of MRPP-substrate complex

The MTO and STO kinetic data suggest the formation of a MRPP1/2/3 complex that enhances catalytic function. To further investigate the formation of a MRPP1/2/3 complex, we performed *in vitro* magnetic capture experiments using nickel (II)-coated magneto-beads (Figure 3). If a stable MRPP1/2/3 complex forms, MRPP3 will elute from the resin along with His_6_-MRPP1/2. Under near-physiological salt conditions (100 mM NaCl), a MRPP1/2/3 complex was not observed, consistent with previous observations (Supplementary Figure S1) (13,29). Even at high MRPP1/2 and MRPP3 concentrations (2.1 μM), the His_6_-MRPP1/2/3 complex is not observed, indicating a weak affinity of MRPP3 for His_6_-MRPP1/2 in the absence of pre-tRNA (Lane 3). Upon addition of pre-tRNA to a reaction containing His_6_-MRPP1/2 and MRPP3 (in CaCl_2_ to inhibit cleavage), MRPP3 co-elutes with His_6_-MRPP1/2 demonstrating the formation of a stable MRPP1/2/3 complex in the presence of pre-tRNA (Lane 6). A control experiment demonstrates that MRPP3 does not elute in the absence of His_6_-MRPP1/2 when pre-tRNA is present (Lane 12). These data reveal an RNA-mediated MRPP1/2/3•pre-tRNA quaternary complex consistent with a recent report by Oerum et al (29).

**Figure 3.**
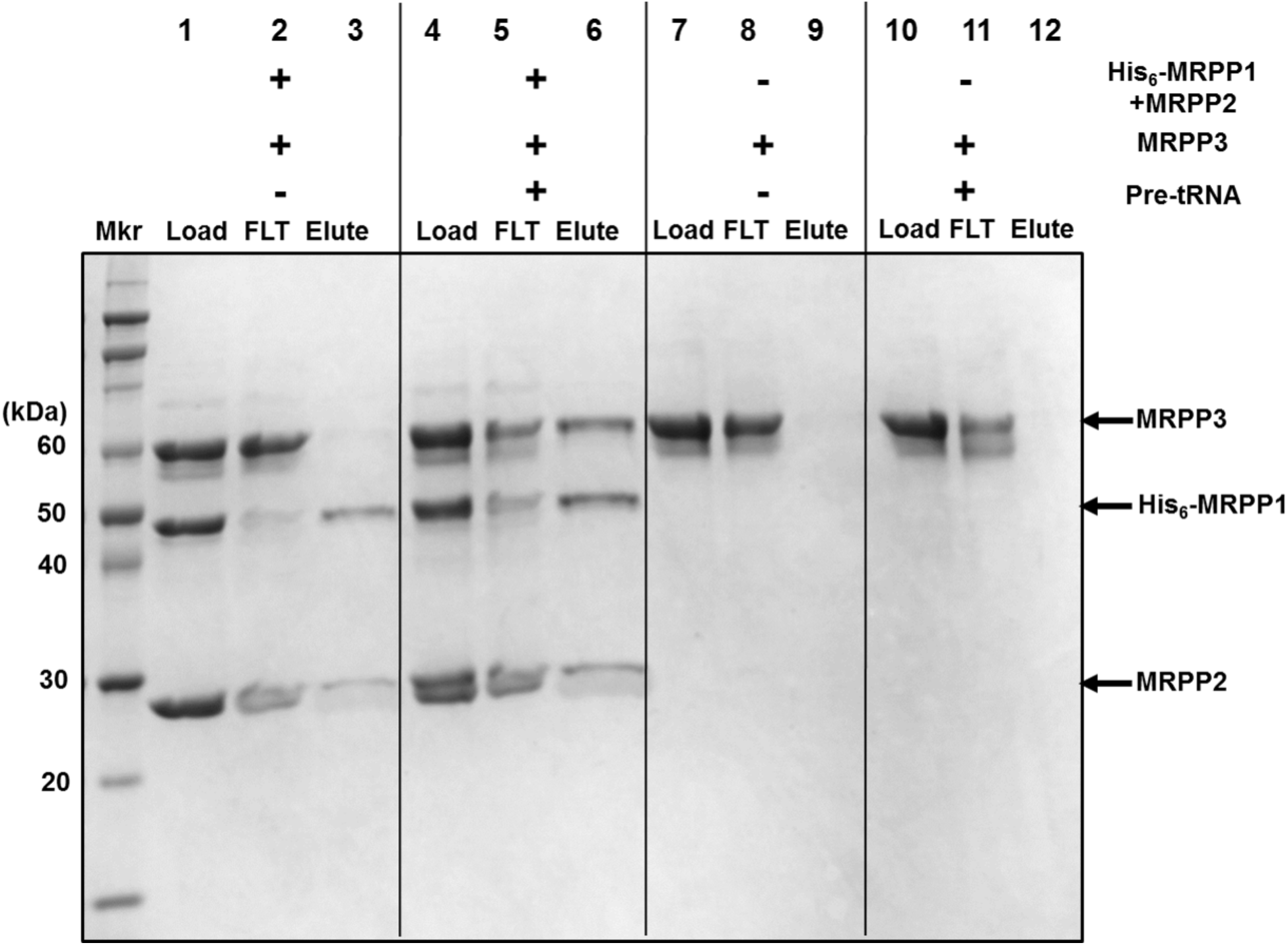
MRPP1/2/3 complex forms in the presence of pre-tRNA. 12% SDS-PAGE stained with Coomassie Blue shows results from magnetic capture experiments performed with 2.1 μM MRPP3 in the absence of pre-tRNA (Lane 1–3, 7–9) or in the presence of 2.1 µM pre-tRNA (Lane 4–6, 10–12) and with the His_6_-MRPP1/2 complex (Lane 1– 6) or without the His_6_-MRPP1/2 complex (Lane 7–12). MRPP3 controls with or without pre-tRNA are in Lanes 7–9 and 10–12, respectively. Samples (Lanes 1, 4, 7, and 10) were loaded onto Ni-NTA magnetic agarose beads and incubated at room temperature for 35 min. The flow-through (FLT) was collected (Lanes 2, 5, 8, and 11). 100 mM EDTA (pH 8) was used to elute the protein and RNA from the beads (Lanes 3, 6, 9, and 12).

### Evidence of synergistic binding between MRPP subunits and pre-tRNA

To evaluate whether MRPP1/2 enhances catalytic activity of RNase P by increasing the pre-tRNA binding affinity of MRPP3, we measured apparent dissociation constants (*K*_D,app_) for each of the individual subunits using a previously described fluorescence anisotropy (FA) binding assay (12,30). We also measured the binding affinity in CaCl_2_, which allows MRPP1/2/3 to bind, but not cleave substrate on the time-scale of our experiments (data not shown) (12,27). In the presence of MgCl_2_ or CaCl_2_, MRPP3-alone weakly binds Fl-pre-tRNA (*K*_D,app_ = 1800 ± 200 nM in MgCl_2_ or *K*_D,app_ = 2300 ± 200 nM CaCl_2_); this value is 1,000-fold weaker than the affinity of the homologous *A. thaliana* PRORP1 for the same substrate (*K*_D,app_ = 1.8 nM in CaCl_2_, Figure 4, Table 2). MRPP1/2 binds Fl-pre-tRNA more tightly (*K*_D,app_ = 40 ± 4 nM) in MgCl_2_ (Figure 4, Table 2). Under CaCl_2_ conditions, the affinity of MRPP1/2 for Fl-pre-tRNA decreases by 30-fold (*K*_D,app_ = 1200 ± 100 nM), Figure 4A, Table 2). Importantly, the addition of MRPP1/2 (600 nM) increases the apparent binding affinity of MRPP3 for Fl-pre-tRNA by nearly 300-fold in CaCl_2_ (*K*_D,app_ = 7 ± 2 nM, Figure 4B, Table 2). No significant change in the FA signal was observed upon addition of MRPP2 alone to Fl-pre-tRNA, consistent with a previous gel shift assay study indicating that MRPP2 does not bind pre-tRNA (23). These data demonstrate that the subunits in the MRPP1/2/3 complex synergize to enhance the substrate binding affinity to a value similar to that of *A. thaliana* PRORP1.

**Table 2.** Dissociation constants (*K*_D,app_) of Fl-pre-tRNA for MRPP proteins.

| | $K_{D,app}$ (nM) | |
| --- | --- | --- |
|  | CaCl<sub>2</sub> | MgCl<sub>2</sub> |
| MRPP1/2 | 1200 ± 100 | 40 ± 4<sup>a</sup> |
| MRPP3 | 2300 ± 200 | 1800 ± 200<sup>a</sup> |
| MRPP3 plus MRPP1/2<sup>b</sup> | 7 ± 2 | NA |
| PRORP1<sup>c</sup> | 1.8 ± 0.3 | NA |

Experiments were done as described in Figure 3. The reported errors are the standard error of the mean obtained from the fit of the FA data.
<sup>a</sup> Experiments were in the presence of MgCl<sub>2</sub>, which does not activate pre-tRNA cleavage when MRPP1/2 or MRPP3 are present alone.
<sup>b</sup> MRPP1/2 concentration is 600 nM.
<sup>c</sup> The binding affinity of PRORP1 was measured with 2 nM Fl-pre-tRNA and a quadratic equation (Equation 11) was used to fit the data.
NA: binding measurement not available because cleavage occurs in MgCl<sub>2</sub>.

**Figure 4.**
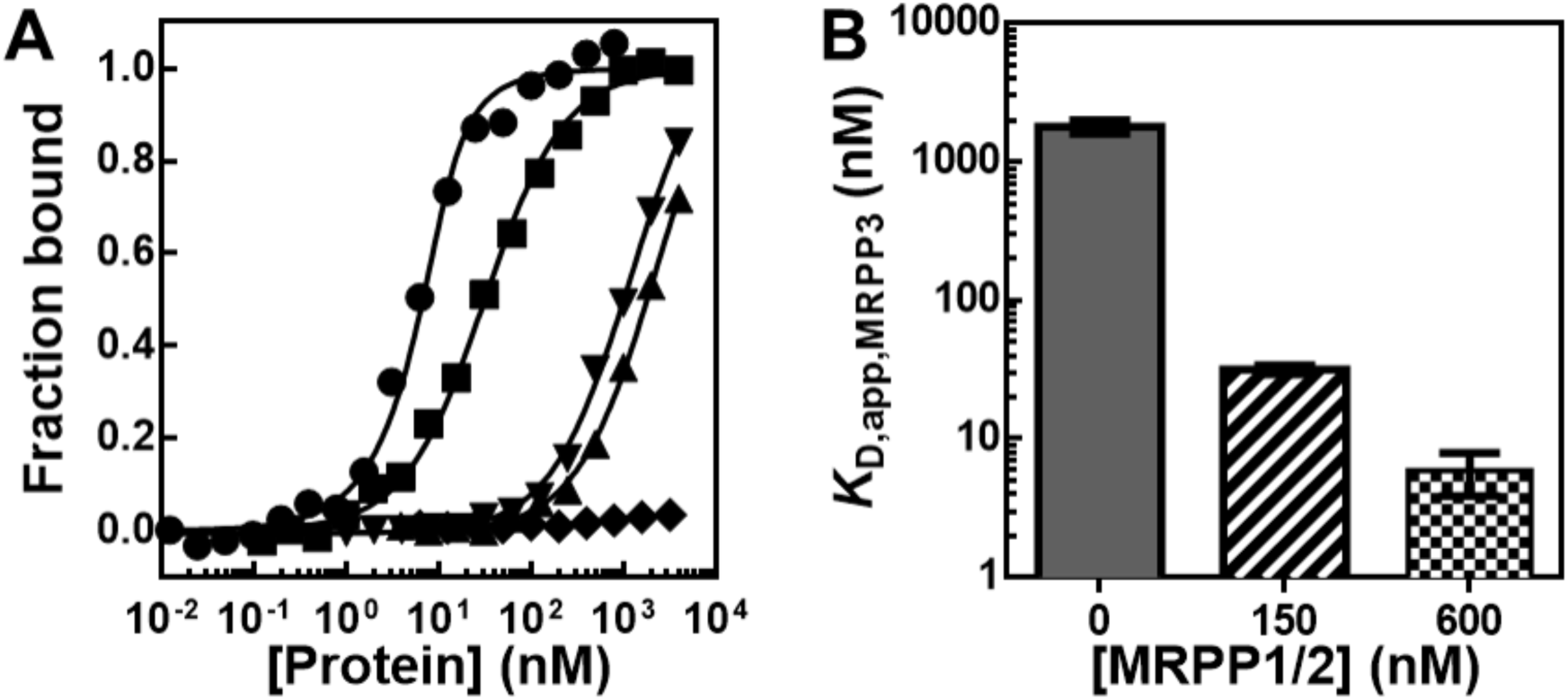
Binding affinity of mitochondrial RNase P proteins to Fl-pre-tRNA in the presence of CaCl_2_. (**A**) Fraction bound calculated from the FA binding assay (Equation 9) measuring the binding of 10 nM Fl-pre-tRNA to varied concentrations of MRPP1/2 (inverted triangles), MRPP3 (triangles), MRPP2 (diamonds), and MRPP3 with 150 nM MRPP1/2 (squares). The concentration of Fl-pre-tRNA was reduced to 2 nM for PRORP1 (dots). FA is measured after a 2-5 min incubation. A binding isotherm (Equation 10) was fit to the data except for PRORP1, which was fit to a quadratic equation (Equation 11). Dissociation constants (*K*_D,app_) are listed in Table 2. (**B**) Apparent affinity of MRPP3 (*K*_D,app,MRPP3_) to Fl-pre-tRNA in the presence of MRPP1/2. The FA binding assay was performed using 10 nM Fl-pre-tRNA with varying MRPP3 concentrations at a constant MRPP1/2 concentration (0, 150, or 600 nM). The errors reported are from the fit of the fraction bound curve obtained from FA data (11 to 17 protein concentrations) as shown in (A).

### MRPP3 recognizes MRPP1/2-pre-tRNA as co-substrate

MRPP1/2 functions similarly to a co-substrate for mtRNase P catalysis. Therefore, we hypothesized that the kinetic stoichiometry measured in reactions (Figure 2) does not necessary equal to a fixed thermodynamic stoichiometry of MRPP1/2/3 ternary complex. We measured the ratio of MRPP1/2 to MRPP3 in the ternary complex from the activation of the MRPP3 cleavage activity by MRPP1/2 under STO conditions. As described above, both the values of *k*_cat,app_ (MTO) and *k*_obs_ (STO) catalyzed by MRPP3 display a hyperbolic dependence on the concentration of MRPP1/2 with *K*_1/2,app_^MRPP1/2^ values in the low µM range (Figure 1D and Figure 2B). The maximal STO activity (*k*_max,app_) depends on the concentrations of both MRPP1/2 and MRPP3 (Figure 2B and 2C). These data are consistent with measurement of apparent binding constants involving multiple steps, rather than a single stoichiometric formation of a MRPP1/2/3 complex. Consistent with our hypothesis, the ratio of MRPP1/2 to MRPP3 needed to obtain maximal STO activity (*k*_max,app_) is variable (Figure 2B and 2C). These data show that the stoichiometry of MRPP subunits (MRPP1:2:3) to achieve the maximal activity is not a simple 2:4:1 ratio, as commonly concluded in previous reports (13,14,18,24-26).

### The MRPP1/2 subcomplex does not enhance the apparent metal affinity of MRPP3

A possible role for MRPP1/2, as proposed by recent structural studies of MRPP3 (25,26), is to enhance binding of metal ions in the active site. Both reported MRPP3 crystal structures lack large portions of the N-terminal region and appear to be in an inactive conformation with no metal ions bound to the active site (25,26). To test whether MRPP1/2 activates MRPP3 catalysis by positioning catalytic metal ions in the active site, we measured activation of cleavage activity of MRPP3 by Mg^2+^ (*K*_1/2_^Mg^) at multiple concentrations of MRPP1/2 under MTO conditions (Figure S2, Equation S1). If MRPP1/2 enhances the affinity of the active site for Mg^2+^, we predict that the value of *K*_1/2_^Mg^ would decrease with increasing MRPP1/2 concentration. However, *K*_1/2_^Mg^ remains unchanged between 150 nM and 500 nM MRPP1/2 (*K*_1/2_^Mg^ = 1.9 ± 0.2 mM and 2.5 ± 0.6 mM, respectively). Thus, our data do not support the hypothesis that the MRPP1/2 subcomplex enhances the affinity of catalytic Mg^2+^ ions to MRPP3.

## DISCUSSION

### Kinetic model for mitochondrial RNase P

We sought to understand the role of MRPP1/2 in human MRPP1/2/3 activity. The simplest model for pre-tRNA cleavage catalyzed by human mtRNase P involves the formation of a tertiary protein complex (MRPP1/2/3), followed by the binding and cleavage of pre-tRNA (Scheme 1). However, the data presented in this work and others (13,18,25,26,29) show that MRPP3 does not strongly associate with MRPP1/2 under physiological conditions (13) and pre-tRNA binds to both MRPP1/2 and MRPP3 subunits (Figure 4, Table 2). Furthermore, synergistic binding between MRPP1/2/3 and pre-tRNA is observed (Table 2) and could possibly be explained by the stabilization of the MRPP1/2/3 complex by pre-tRNA (Figure 3). This minimal model also predicts that MTO activity at saturating pre-tRNA is not dependent on the concentration of MRPP1/2 (*k*_cat,app_ = *k*_chem_, see derivation for Scheme 1 and Equations S2, S3 in the Supplementary Data), but this is not true in our experiments where a hyperbolic dependence is clearly observed (Figure 1D).

A kinetic model that aligns with both our MTO and binding measurements is shown in Scheme 2. In this model, pre-tRNA binds to MRPP1/2, MRPP3 or the MRPP1/2/3 complex to form catalytically active MRPP1/2/3•pre-tRNA. The Equations describing kinetic parameters for Scheme 2 explain the hyperbolic dependence on the MRPP1/2 concentration observed for *k*_cat,app_ and *K*_M,app_ (Figure 1; Scheme 2, Supplementary Equations S5, S6, and S7, respectively). Furthermore, no inhibition of the STO rate constant was observed at high concentrations of either MRPP3 or MRPP1/2 (Figure 2), indicating that both MRPP3•pre-tRNA and MRPP1/2•pre-tRNA complexes are productive and on a pathway to form a catalytically active MRPP1/2/3•pre-tRNA complex. Therefore, we propose a mechanism with a closed thermodynamic box shown in Scheme 2.

Our experimental data also allow estimation of values for the equilibrium constants shown in Scheme 2: *K*_1_ reflects the measured dissociation constant between pre-tRNA and MRPP1/2 (0.04 μM, Table 2); *K*_3_ is the measured dissociation constant for pre-tRNA binding to MRPP3 (1.8 μM, Figure 3 and Table 2); *K*_4_ is approximated by the measured *K*_1/2_ for enhancement by MRPP1/2 of either the MTO *k*_cat,app_ (0.4 ± 0.1 μM) (Figure 1D) or the STO *k*_max,app_ (0.3 ± 0.2 μM, Figure S4); and *K*_6_ is estimated from the *K*_1/2_ for activation of activity by MRPP3 at saturating MRPP1/2 under STO conditions (0.09 μM, Figure 2C). The value for *K*_5_, which describes the affinity of pre-tRNA for the MRPP1/2/3 tertiary complex, is estimated as ≤0.007 μM based on binding experiments (Figure 3, Table 2). However the thermodynamic box in Scheme 2 requires that *K*_1_*K*_6_ = *K*_3_*K*_4_ = *K*_2_*K*_5_ (Equation S4), which is not the case based on our data (*K*_3_*=* 1.8 µM and *K*_4_ = 0.4 µM, therefore *K*_3_*K*_4_ is comparable within error to *K*_2_*K*_5_ = 0.8 ± 0.1 µM^2^, but it is not equal to *K*_1_*K*_6_ since *K*_1_*=* 0.04 µM and *K*_6_ = 0.09 µM).

Since the above calculated thermodynamic box is not balanced, the kinetic mechanism may include additional steps that are not shown in Scheme 2. Therefore, we propose a modified kinetic model that includes one additional step (*K*_7_): a conformational change step following MRPP1/2 binding to pre-tRNA (Scheme 3). We propose that *K*_1,app_ is the apparent binding affinity of pre-tRNA for MRPP1/2 (0.04 μM, Table 2, Equation 5 and 6) and reflects both the binding and conformational change steps (*K*_1_). The values for *K*_1_ (∼0.04 μM) and *K*_7_ (∼0.005) were estimated from *K*_1,app_ using Equation 5. The value of *K*_7_ (∼0.005) indicate that this conformational change step is unspontaneous (ΔG>0). There is precedence for unspontaneous conformational change steps utilized by multiple important enzymes for ligand recognition (31,32) and a conformational change step has been observed for *B. subtilis* RNase P catalysis (33) and in PRORP2 structures (34).

Scheme 3 is also consistent with the results from pull-down and MTO experiments. The estimated binding affinity between MRPP1/2 and MRPP3 (*K*_2_) is very weak at ≥103 μM (Equation 6), which could explain why no MRPP1/2/3 complex is observed at 2.1 μM of each MRPP proteins (Figure 3). Although Scheme 3 predicts a hyperbolic dependence of (*k*_cat_/*K*_M_)_app_ on the concentration of MRPP1/2 (Equation 3), the observed linear dependence of (*k*_cat_/*K*_M_)_app_ on the concentration of MRPP1/2 (Figure 1F) is consistent with an linear approximation that can be derived from Equation 3 where the experimental concentration of MRPP1/2 (low μM) is much lower than *K*_2_ (103 μM). In addition, we derived Michaelis-Menten equations for many possible thermodynamic models. All kinetic models that did not contain at least one conformational change step did not fit with all of our data. Therefore, we present Scheme 3 as a simplest model that will fit all experimental data and the requirements of a thermodynamic box (Equation 4, 5 and 6).

In summary, Scheme 3 best fits our experimental data, and both qualitatively and quantitatively illustrates the intricate mechanism employed by MRPP1/2 and MRPP3 in mtRNase P catalysis. We propose that the role of MRPP1/2 is to increase the binding affinity of pre-tRNA either by directly contacting pre-tRNA or by altering the structure of MRPP3 or pre-tRNA in a MRPP1/2/3•pre-tRNA complex. Additionally, MRPP1/2 significantly increases the catalytic activity likely by participating in protein-protein and/or protein-RNA contacts, interacting with pre-tRNA and/or MRPP3.

### Interplay of pre-tRNA and the MRPP1/2/3 complex

Our data are consistent with structural studies reporting the MRPP1/2/3-tRNA complex and a distorted MRPP3 active site (25,26,29) and provide insights into the kinetics and thermodynamics of these interactions. Pre-tRNA stabilizes the MRPP1/2/3 complex as demonstrated by magnetic capture (Figure 3) and binding data (Figure 4, Table 2). Furthermore, pre-tRNA has a greater affinity for the full MRPP1/2/3 complex than the separate subunits (Scheme 3). We speculate that upon binding pre-tRNA, changes in protein conformations may lead to changes in protein-protein contacts between MRPP1/2 and MRPP3, as previously proposed (26,29); and/or alteration of the pre-tRNA structure which leads to enhanced recognition by the catalytic subunit in MRPP3.

The enhanced catalytic efficiency of MRPP1/2/3 may be due to rearrangements in the MRPP3 active site or the bound pre-tRNA structure. MRPP1/2 may play a role in selecting substrates. Given that mammalian mitochondrial tRNAs exhibit noncanonical features, this may provide an explanation for evolving a multi-protein complex for human mtRNase P (5,35,36).

### Roles of MRPP1/2 in mitochondrial pre-tRNA processing

The MRPP1/2 subcomplex has a profound effect on pre-tRNA processing, as shown by a significant decrease in catalytic efficiency (100,000-fold, Table 1) and substrate affinity (300-fold, Table 2) when levels of MRPP1/2 are reduced. These results corroborate *in vivo* data where siRNA knockdowns of MRPP1 and MRPP3 show differential effects on pre-tRNA processing (6). When MRPP1 and MRPP3 are reduced by similar levels, the MRPP1 knockdown results in a 10-fold higher accumulation of some pre-tRNA transcripts compared to the MRPP3 knockdown. These *in vivo* observations combined with our *in vitro* kinetic data (Figure 1) highlight the obligate role of MRPP1/2 in human mtRNase P catalysis. MRPP1/2 may facilitate catalysis by altering the contacts between MRPP3 and pre-tRNA, stabilizing a catalytically competent structure (Scheme 3). Future work such as transient kinetic characterizations are needed to dissect this mechanism.

In conclusion, MRPP1/2 plays an essential role in mtRNase P by enhancing the catalytic efficiency and substrate affinity of MRPP3. Our proposed model (Scheme 3) provides a framework for investigating and identifying the detailed steps leading to mtRNase P activity. This finding reveals a mechanism likely underpinning the roles of MRPP1/2 in multiple mt-tRNA processing pathways.

## EXPERIMENTAL PROCEDURES

### Pre-tRNA preparation

Fluorescein-labeled *B. subtilis* pre-tRNA^Asp^ (Fl-pre-tRNA) was prepared as previously described with a 5-nucleotide leader and a discriminator base at the 3’ end without the 3’ CCA sequence (27). The template was produced from plasmid DNA containing a T7 promoter region and the *B. subtilis* pre-tRNA^Asp^ sequence by PCR amplification. The *in vitro* transcription reaction was performed using the PCR template in the presence of guanosine 5’-monothiophosphate (GMPS) (37,38). 5’-GMPS-pre-tRNA^Asp^ was prepared by *in vitro* transcription in the presence of guanosine 5’-monothiophosphate (GMPS) and labeled by reaction with 5-iodoacetamido-fluorescein (5-IAF, Life Technologies) overnight at 37 °C to prepare Fl-pre-tRNA (37,38). The RNA was purified using a 10% polyacrylamide/bis (39:1) denaturing gel containing 7 M urea. Unlabeled pre-tRNA was prepared using *in vitro* transcription without the addition of GMPS. Nucleoside triphosphates (NTPs) and chemicals were purchased from Sigma. Inorganic pyrophosphatase was purchased from Roche Applied Science. GMPS was synthesized from 2’, 3’ isopropylideneguanosine and thiophosphoryl chloride (39). Recombinant His_6_-T7 RNA polymerase was expressed in *E. coli* and purified by Ni-NTA chromatography (40). Before each assay, FL-pre-tRNA was unfolded at 95 °C for 2 min in water, and refolded by incubation at 37 °C for 15 min followed by addition of the desired buffer and incubation for another 30 min at 37 °C.

### Purification of recombinant human MRPPs

To prepare a plasmid for the co-expression and co-purification of the MRPP1/2 subcomplex to achieve increased yield and stability of recombinant MRPP1, DNA encoding an N-terminal truncation of the MRPP1 gene (TRMT10C) (residues 40-403, Δ39MRPP1) with an N-terminal His6-tag followed by a TEV protease site and the full-length MRPP2 gene (SDR5C1) were cloned into a pCDFDuet-1 (Novagen) co-expression vector as previously described (41). The plasmid was transformed and expressed in Rosetta (DE3) or Rosetta2 (DE3) E. coli cells. Cells were grown in Terrific Broth or auto-induction media (Novagen) containing 50 mg/L streptomycin and 34 mg/L chloramphenicol. Cells grown in Terrific Broth were grown to an OD600 of 0.6-0.8 at 37 °C, and then induced by addition of 660 µM β-D-1-thiogalactopyranoside (IPTG) followed by growth for 16 h at 18°C. Cells grown in auto-induction media were incubated for 24 h at 37 °C. Cells were harvested and lysed using a microfluidizer in buffer A (20 mM Tris-HCl or 50 mM 3-(N-Morpholino)propanesulfonic acid (MOPS) (pH 7.5), 1 mM Tris(2-carboxyethyl)phosphine (TCEP), and 10% glycerol) containing 1 M NaCl, 15 mM imidazole and one tablet of Complete EDTA free protease inhibitor cocktail (Roche). Cell lysate was pelleted for 60 min at 30,600 × g at 4 °C. The soluble fraction was then applied to a Ni-Sepharose column preequilibrated with buffer A containing 150 mM NaCl. The column was washed with over 10 column volumes (CV) with buffer A containing 150 mM NaCl followed by 1 M NaCl (in buffer A) wash. Bound proteins were eluted by gradient from 50-500 mM imidazole in buffer A over 10 CV. Fractions containing MRPP1/2 were pooled, His_6_TEV protease was added, and the reaction was dialyzed against buffer A containing 150 mM NaCl. The sample was then applied to a second Ni-Sepharose column. The flow-through was collected, concentrated, and dialyzed into 50 mM MOPS (pH 7.5), 300 mM NaCl, and 1 mM TCEP. The sample was loaded onto a Sephacryl-200 gel-filtration column (GE Healthcare) preequilibrated with 50 mM MOPS, 1 mM TCEP, and 300 mM NaCl. Protein was eluted at 1 mL/min and the A280 was monitored. Peak fractions were analyzed by SDS-PAGE. Fractions containing MRPP1 and MRPP2 were pooled. The molecular extinction coefficients (ε_280_) of the individual subunits are 62,130 M-1cm-1 and 4,720 M-1cm-1 for Δ39MRPP1 and MRPP2, respectively. The ratio of MRPP1/2 were determined to be 2:4 for calculation of concentration of MRPP1/2 complex as detailed in (41).

The MRPP3 gene (*KIAA0391*) with an N-terminal truncation (residues 46-583, Δ45MRPP3) was cloned into a pETM-11 vector that adds an N-terminal His_6_-tag as previously described (42). The plasmid was transformed and expressed in Rosetta (DE3) *E. coli* cells. Cells were grown in LB medium at 37 °C to an OD_600_ of 0.6-0.8, and then induced by addition of 660 µM IPTG followed by growth for 16 h at 18°C. Cells were lysed using a microfluidizer and centrifuged for 60 min at 30,000 × g at 4 °C. The resulting soluble fraction was then applied to a Ni-Sepharose column (GE Healthcare Life Sciences) preequilibrated in buffer B (50 mM MOPS, pH 7.5, 10% glycerol, 150 mM NaCl, 1 mM MgCl_2_, 1 mM TCEP) with 15 mM imidazole. Buffer B containing 1 M NaCl was used to elute nucleic acid contaminants. Bound proteins were eluted using an imidazole gradient (50-500 mM) in buffer B over 10 CV. Fractions containing MRPP3 were pooled and TEV protease was added and the sample was dialyzed overnight against Buffer B at 4 °C. The sample was then applied to a second Ni-Sepharose column. The flow-through was collected, concentrated, and dialyzed into 50 mM MOPS (pH 7.5), 1 mM TCEP, 150 mM NaCl, and 10 % glycerol. Full-length MRPP2 was expressed in *E. coli* and purified in a similar manner to that for Δ45MRPP3 as described in (42). The concentrations of Δ45MRPP3 and MRPP2 were determined by absorbance (Δ45MRPP3, ε_280_ = 85,830 M^−1^cm^−1^; MRPP2, ε_280_ = 4720 M^−1^cm^−1^). PRORP1 was purified as previously described (12).

### Multiple-turnover (MTO) cleavage measurements

MTO assays were performed at limiting concentrations of MRPP3 (25–50 nM) and varied concentrations of MRPP1/2 (150-1200 nM) and Fl-pre-tRNA substrate (150-5000 nM total pre-tRNA containing 40 nM Fl-pre-tRNA). Reactions were initiated by mixing Fl-pre-tRNA with MRPP proteins at 37 °C in reaction buffer (50 mM MOPS pH 7.8, 100 mM NaCl, 4.5 mM MgCl_2_ and 1 mM TCEP). Reaction time points were quenched with an equal volume of 8 M urea, 100 mM EDTA pH 8, 0.05% bromophenol blue, and 0.05% xylene cyanol. Quenched reactions were resolved on a 22% PAGE containing 7 M urea, and imaged by a Typhoon scanner (λ_ex_ = 488 nm, λ_em_ = 535 nm) (Figure 1A). Gels were analyzed using ImageJ software. Initial velocities were determined by 5-12 time points at the linear rate range (Figure 1B).

The apparent kinetic parameters (*k*_cat,app_, *K*_M,app_, and (*k*_cat_*/K*_M_)_app_) were determined from a hyperbolic fit (43) of the dependence of the initial velocities divided by the MRPP3 concentration on the MRPP1/2 concentration. The steady-state kinetic equations were derived for *k*_cat,app_ (Equation 1), *K*_M_,_app_ (Equation 2), and (*k*_cat_*/K*_M_)_app_ (Equation 3) from the model in Scheme 3 (Equation 4) (see derivations in Supplementary Data). Equation 1 was fit to the MRPP1/2 dependence of *k*_cat,app_ to determine the value of *k*_chem_ at saturating MRPP1/2 concentration and the value for *K*_4_ (Figure 1D). Equation 2 was fit to the MRPP1/2 dependence of *K*_M,app_ to estimate values for *K*_4_ and *K*_2_*K*_5_ (Figure 1E). In this analysis, the error in the *K*_2_*K*_5_ value is large (>100%). The value of *K*_M,app_ approaches *K*_5_ at saturating MRPP1/2 (Equation 2). Equation 3 predicts a linear dependence of (*k*_cat_*/K*_M_)_app_ on MRPP1/2 under these experimental conditions ([M1/2]<<*K*_2_). Thus, the value of the linear slope (Equation 3, Figure 1F) and the value of *k*_chem_ (Equation 1) were used to calculate the value of *K*_2_*K*_5_. (within 2-fold of the value of *K*_2_*K*_5_ estimated using Equation 2) which is used in subsequent analyses. The values of *K*_1,app_ and *K*_3_ were measured by fluorescence anisotropy (FA) binding assays (see below). The values for *K*_1_ and *K*_7_ were calculated using *K*_1,app_ and Equations 5 and 6.

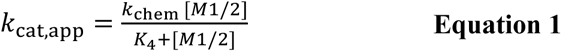

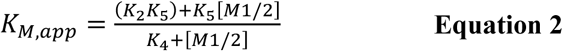

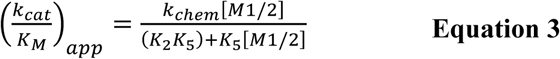

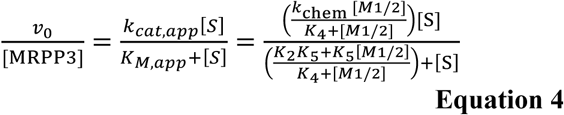

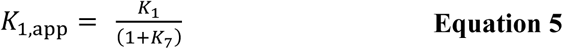

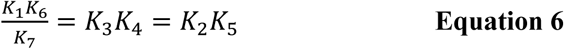

### Single-turnover (STO) experiments

STO activity was measured using 10 nM Fl-pre-tRNA and 0.1 –10 µM MRPP3 pre-incubated with 0.2–1.6 µM MRPP1/2 subcomplex in 50 mM MOPS (pH 7.8), 150 mM NaCl, 4.5 mM MgCl_2_, 2 mM TCEP at 37 °C. Time points were taken and analyzed by urea-PAGE as described for MTO experiments. A single exponential equation (Equation 7) was fit to the percent cleavage (*Y*_*c*_) time course to calculate the single-turnover rate constant (*k*_obs_). Equation 8 was first fit to the dependence of *k*_obs_ (represented as A) on MRPP1/2 concentration (shown as [P]) to calculate the apparent affinity of MRPP1/2 (*K*_1/2_^MRPP1/2^, represented as C) and *k*_max,app_ (represented as B). This was repeated for different MRPP3 concentrations (Figure 2B). Equation 8 was also fit to the dependence of *k*_max,app_ on MRPP3 concentration to calculate the apparent affinity of MRPP3 (C = *K*_1/2_^MRPP3^), which is approximately equal to the dissociation constant of MRPP3 from the MRPP1/2-bound substrate (*K*_6_) under saturating MRPP1/2 conditions (Figure 2C).

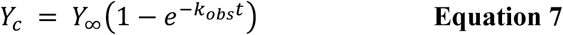

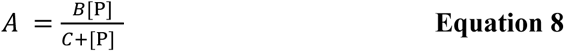

### Magnetic capture assay

We applied magnetic capture methods (44) using a His_6_-MRPP1/2 subcomplex (His_6_-tagged MRRP1 in complex with tag-less MRPP2) to pull-down MRPP3 *in vitro*. The Ni-NTA magnetic beads (Qiagen) were pre-equilibrated in binding buffer (50mM HEPES pH 7.8, 100 mM NaCl, 4.5 mM CaCl_2_, and 1 mM TCEP) before incubation with samples containing 2.1 µM His_6_-MRPP1/2 subcomplex and 2.1 µM MRPP3 with or without 2.1 µM pre-tRNA. The Ni-NTA magnetic beads were incubated with samples for 35 min at room temperature before the flow-through was collected and the beads were washed with binding buffer twice. The samples were eluted with 100 mM EDTA (pH 7.5), and were then run on a SDS-PAGE gel stained with Coomassie Blue.

### Fluorescence anisotropy (FA) binding assay

Binding experiments using 10 nM Fl-pre-tRNA and varying concentrations of proteins were carried out in 50 mM MOPS (pH 7.8), 100 mM NaCl, 1 mM TCEP, and 4.5 mM MgCl_2_ (physiologically relevant divalent metal ion) or CaCl_2_ (divalent ion in which cleavage by MRPP1/2/3 is inhibited to measure substrate binding). Proteins and Fl-pre-tRNA were incubated at 37 °C for 2 min and the FA values measured in a 384-well plate using a TECAN plate reader. The FA binding data were converted to fraction bound (*Y*_*b*_) using Equation 9, where *A*_*0*_ is the FA value of unbound Fl-pre-tRNA and *A*_*b*_ is the FA value of fully bound protein-Fl-pre-tRNA complex. All fraction bound data, with the exception of PRORP1, were fit to a binding isotherm (Equation 10, where [P] is the protein concentration). PRORP1 data were fit to a quadratic equation (Equation 11 where *L* is the concentration (2 nM) of the Fl-pre-tRNA) because the Fl-pre-tRNA concentration is close to the value of the dissociation constant (*K*_D_).

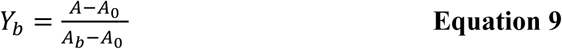

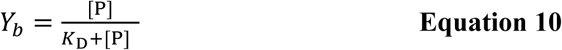

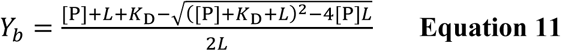

## Acknowledgements

The authors thank Dr. Jane Jackman (Ohio State University) for the kind gift of a MRPP1 plasmid and discussions. The authors also thank Dr. Agnes Karasik, Andrea Stoddard and Dr. Elaina Zverina for helpful suggestions regarding the manuscript, molecular biology and protein purification strategies.

## Conflict of interests

The authors declare that they have no conflicts of interest with the contents of this article.

## Author contributions

X. L. and N. W. designed and carried out most of the experiments, analyzed the results, and wrote the manuscript. A.S. further optimized the MRPP1/2 protein purification. B. P. K., M. J. H., and W. H. L. carried some of the molecular biology and protein purifications and provided feedbacks to the manuscript. M. K. and C. A. F supervised the project, provided funding, and revised the manuscript.

## FOOTNOTES

This work was supported by the National Institutes of Health [R01 GM117141 to M.K., R01 GM55387 to C.A.F].

The abbreviations used are: mtRNase P, human mitochondrial RNase P; MRPP, mitochondrial RNase P protein; MTO, multiple-turnover; STO, single-turnover; PRORP, protein-only RNase P protein; Fl-pre-tRNA, *B. subtilis* fluorescein-labelled pre-tRNA^Asp^; M1/2, MRPP1/2 complex.

---

**Scheme 1.**
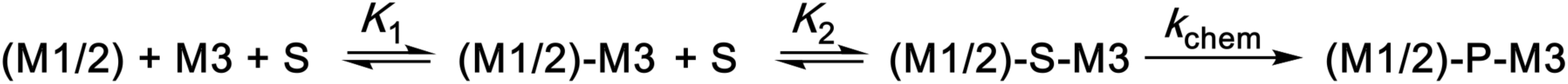
Simple kinetic model for human mtRNase P. MRPP3 is denoted by M3, MRPP1/2 subcomplex is denoted by (M1/2), pre-tRNA substrate is denoted by S, and tRNA and 5’ leader products are denoted by P. (M1/2) binds to M3, after which (M1/2)-M3 binds to S to catalyze cleavage and generate P.

**Scheme 2.**
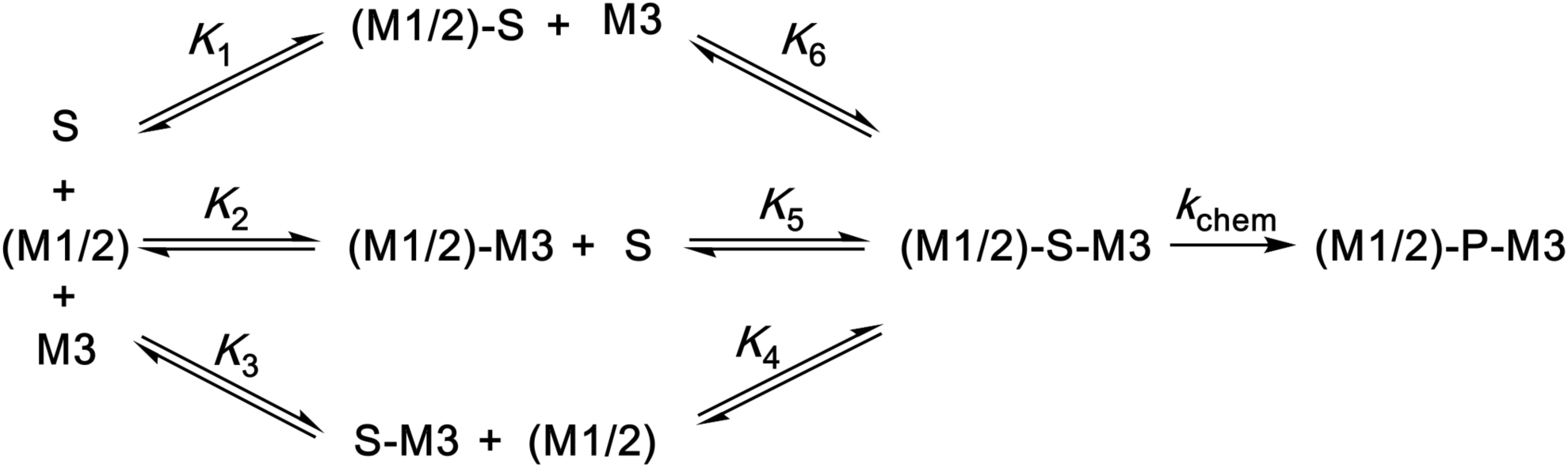
Branched model for human mtRNase P. The (M1/2)-S-M3 ternary complex can be formed through three pathways. In one pathway, S binds to (M1/2) first, followed by binding to M3 (top). In the second pathway, S binds to a (M1/2)-M3 complex to form the (M1/2)-S-M3 ternary complex (center). In the third pathway, S binds to M3 followed by addition of the (M1/2) complex (bottom).

**Scheme 3.**
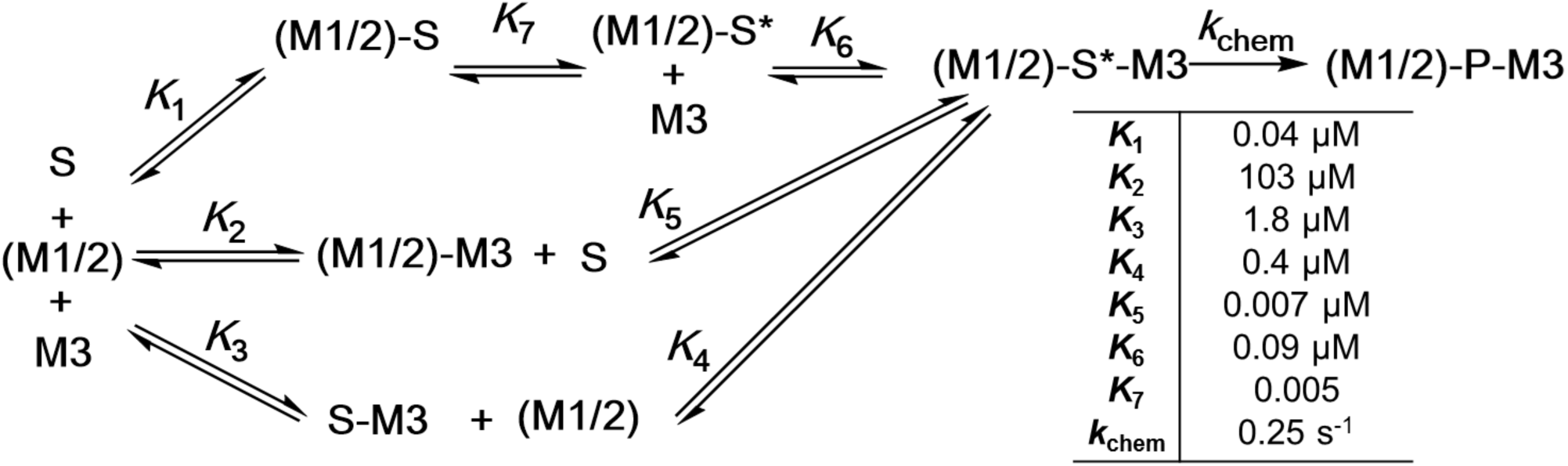
New kinetic model for human mtRNase P. A conformational change step (*K*_7_) is proposed in the first pathway: S binds to (M1/2) through *K*_1_, followed by a conformational change step (*K*_7_) to form (M1/2)-S*, and binding to M3 through *K*_6_ to form an active (M1/2)-S*-M3 complex. The values of the thermodynamic constants and the cleavage rate constant are listed in the table.

## Footnotes

1 This calculation is based on observing <10% product formation on urea-PAGE gel and assuming a mechanism where product release is not rate limiting and the pre-tRNA concentration is sub-saturating.

