## Supplemental Info for "Kinetic mechanism of human mitochondrial RNase P"

#### SUPPLEMENTARY METHODS

##### Magnesium-dependence

The magnesium dependence of cleavage of pre-tRNA by human mitochondrial RNase P was measured under MTO conditions using 500 nM pre-tRNA containing 40 nM Fl-pre-tRNA, sub-saturating (150 nM) or saturating (500 nM) MRPP1/2, in 50 mM MOPS (pH 7.8) and 1 mM TCEP (pH 7.5). The initial velocity was measured over six concentrations of $Mg^{2+}$ holding the ionic strength constant by varying NaCl concentration (μ = 115 mM taking only the concentrations of NaCl and $MgCl_2$ into account). For reactions using 150 nM MRPP1/2, 30 nM MRPP3 was used and for reactions using 500 nM MRPP1/2, 40 nM MRPP3 was used. Equation S1 was fit to the $Mg^{2+}$ dependence of the apparent steady state parameters where $v_o$ is the initial velocity, $V_{max}$ represents the maximal velocity at saturating $Mg^{2+}$, and $K_{1/2}^{Mg}$ is the concentration of $Mg^{2+}$ at which $V_{max}$ is half-maximal.

$$v_o = \frac{V_{max}[Mg^{2+}]}{K_{1/2}^{\mathrm{Mg}} + [Mg^{2+}]} \qquad \textbf{Equation S1}$$

[†] The authors wish it to be known that, in their opinion, the first two authors should be regarded as joint First Authors.

### SUPPLEMENTARY FIGURES

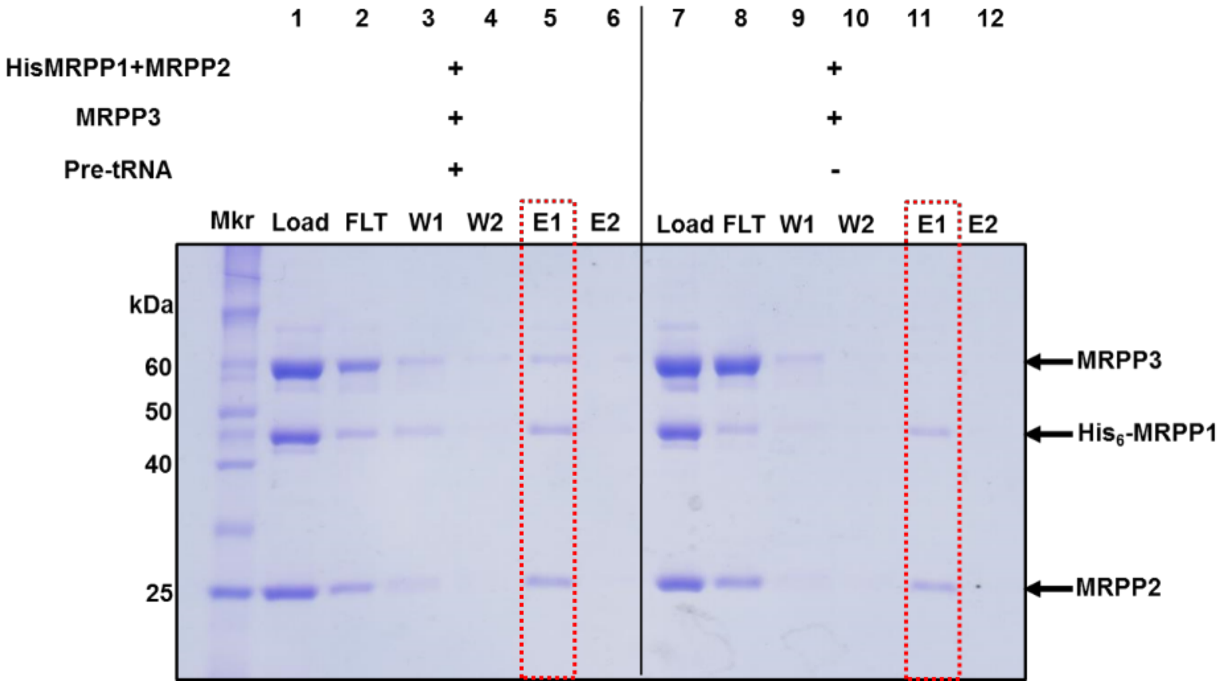


**Figure S1.** MRPP1/2/3/pre-tRNA complex forms in the presence of pre-tRNA. 12% SDS-PAGE stained with Coomassie Blue showing results from pull-down assays performed with MRPP3 in the presence of pre-tRNA (Lane 1–6) or in the absence of pre-tRNA (Lane 7-12). Conditions and procedures similar as described in Figure 2 except for the NaCl concentration is 150 mM. Mkr: protein size marker. FLT: flow-through. W: wash. E: elute.

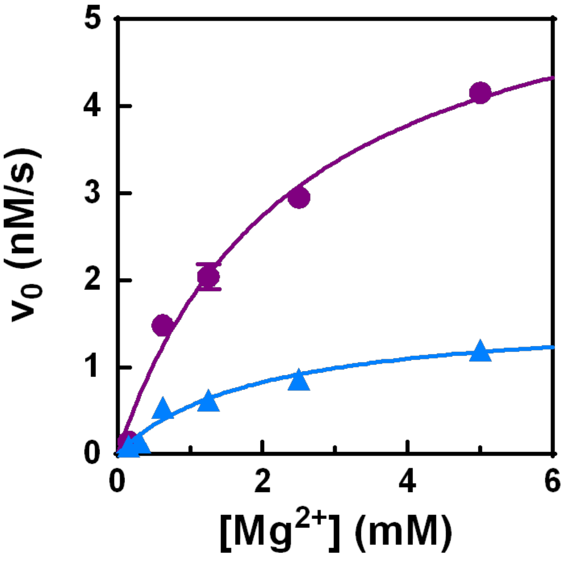

**Figure S2.** Magnesium dependence of the MTO activity of mtRNase P. Initial velocities were measured using the same conditions as in Figure 1 at varying concentrations of $MgCl_2$ and MRPP1/2 (500 nM, purple; 150 nM, blue) with sub-saturating pre-tRNA (500 nM). Equation 6 (main text) was fit to the data to calculate $K_{1/2}^{Mg}$ values of 2.5 ± 0.6 mM and 1.9 ± 0.2 mM for 500 nM and 150 nM MRPP1/2, respectively. The error bars are the standard errors of the mean values obtained from fitting the data.

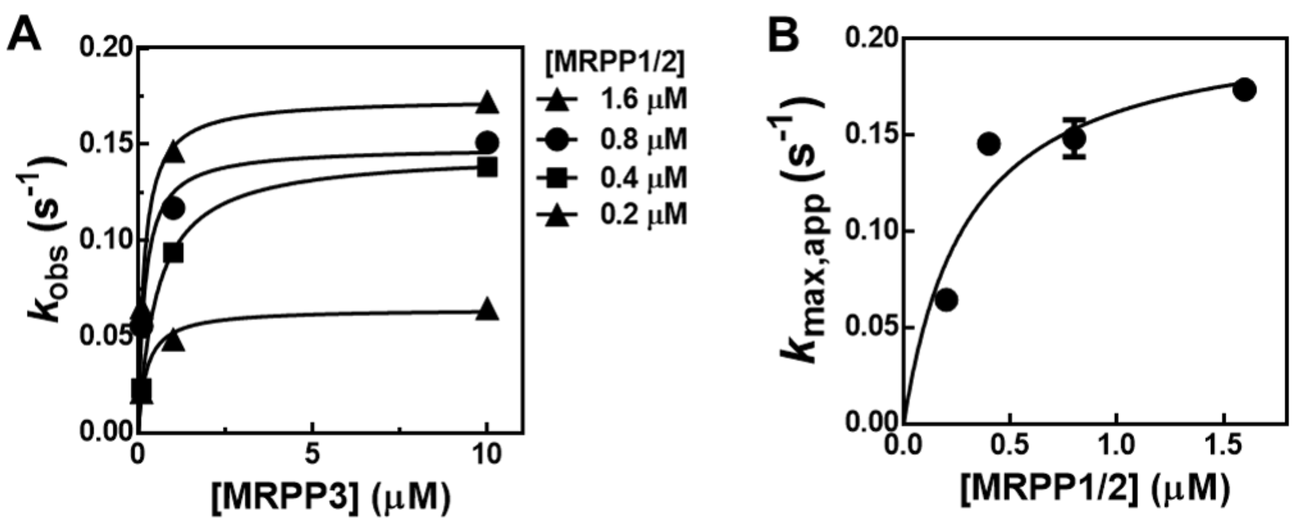


**Figure S3.** Re-plotting the STO data in Figure 2B to estimate $K_4$ and $k_{chem}$. (**A**) Equation 8 (main text) was fit to the MRPP3 dependence of $k_{obs}$ at various concentrations of MRPP1/2 to determine $k_{max,app}$. (**B**) Equation 8 was fit to the MRPP3 dependence of $k_{max,app}$ to determine an apparent $K_{1/2}^{MRPP1/2}$ (0.3 ± 0.2 μM) which is within error of the values of $K_4$ determined from MTO data (Figure 1D), and $k_{max}$ (0.21 ± 0.04 $s^{-1}$) which is within error of the $k_{chem}$ value determined from Figure 1D and Figure 2C.

**DERIVATIONS**

**Derivation of Michaelis–Menten kinetic equations according to Scheme 1**

$$rate = k_{chem}[M1/2 \cdot S \cdot M3]$$

$$[M3_{tot}] = [M3] + [M1/2 \cdot M3] + [M1/2 \cdot S \cdot M3]$$

$$K_1 = \frac{[M1/2][M3]}{[M1/2 \cdot M3]} \; ; \; [M3] = \frac{K_1[M1/2 \cdot M3]}{[M1/2]} = \frac{K_1 K_2[M1/2 \cdot S \cdot M3]}{[M1/2][S]}$$

$$K_2 = \frac{[M1/2 \cdot M3][S]}{[M1/2 \cdot S \cdot M3]} \; ; \; [M1/2 \cdot M3] = \frac{K_2[M1/2 \cdot S \cdot M3]}{[S]}$$

$$[M3_{tot}] = \frac{K_1 K_2[M1/2 \cdot S \cdot M3]}{[M1/2][S]} + \frac{K_2[M1/2 \cdot S \cdot M3]}{[S]} + [M1/2 \cdot S \cdot M3]$$

$$[M3_{tot}] = [M1/2 \cdot S \cdot M3]\left(\frac{K_1 K_2}{[M1/2][S]} + \frac{K_2}{[S]} + 1\right)$$

$$[M3_{tot}] = [M1/2 \cdot S \cdot M3]\left(\frac{K_1 K_2 + K_2[M1/2] + [S][M1/2]}{[M1/2][S]}\right)$$

$$[M1/2 \cdot S \cdot M3] = \frac{[M1/2][M3_{tot}][S]}{K_1 K_2 + K_2[M1/2] + [S][M1/2]}$$

$$[M1/2 \cdot S \cdot M3] = \frac{[M3_{tot}][S]}{\frac{K_1 K_2}{[M1/2]} + K_2 + [S]}$$

$$rate = k_{chem}\left(\frac{[M3_{tot}][S]}{\frac{K_1 K_2}{[M1/2]} + K_2 + [S]}\right)$$ **Equation S2**

$$k_{cat,app} = k_{chem}$$ **Equation S3**

$$K_{M,app} = \frac{K_1 K_2}{[M1/2]} + K_2$$

$$\left(\frac{k_{cat}}{K_M}\right)_{app} = \frac{k_{chem}[M1/2]}{K_1 K_2 + K_2[M1/2]}$$

**Derivation of Michaelis–Menten kinetic equations according to Scheme 2**

$$rate = k_{chem}[M1/2 \cdot S \cdot M3]$$

$$[M3_{tot}] = [M3] + [S \cdot M3] + [M1/2 \cdot M3] + [M1/2 \cdot S \cdot M3]$$

$$K_1 = \frac{[M1/2][S]}{[M1/2 \cdot S]} \; ; \; [M1/2 \cdot S] = \frac{[M1/2][S]}{K_1}$$

$$K_2 = \frac{[M1/2][M3]}{[M1/2 \cdot M3]} \; ; \; [M1/2 \cdot M3] = \frac{[M12][M3]}{K_2}$$

$$K_3 = \frac{[M3][S]}{[S \cdot M3]}$$

$$K_4 = \frac{[M1/2][S \cdot M3]}{[M1/2 \cdot S \cdot M3]}\ ;\ [S \cdot M3] = \frac{K_4[M1/2 \cdot S \cdot M3]}{[M12]}$$

$$K_5 = \frac{[M1/2 \cdot M3][S]}{[M1/2 \cdot S \cdot M3]}\ ;\ [M1/2 \cdot M3] = \frac{K_5[M1/2 \cdot S \cdot M3]}{[S]}$$

$$K_6 = \frac{[M1/2 \cdot S][M3]}{[M1/2 \cdot S \cdot M3]}\ ;\ [M3] = \frac{K_6[M1/2 \cdot S \cdot M3]}{[M1/2 \cdot S]}$$

$$K_1K_6 = K_2K_5 = K_3K_4$$ **Equation S4**

$$[M3_{tot}] = \frac{K_6[M1/2 \cdot S \cdot M3]}{[M1/2 \cdot S]} + \frac{K_4[M1/2 \cdot S \cdot M3]}{[M12]} + \frac{K_5[M1/2 \cdot S \cdot M3]}{[S]} + [M1/2 \cdot S \cdot M3]$$

$$[M3_{tot}] = [M1/2 \cdot S \cdot M3]\left(\frac{K_1K_6}{[M1/2][S]} + \frac{K_4}{[M1/2]} + \frac{K_5}{[S]} + 1\right)$$

$$[M3_{tot}] = [M1/2 \cdot S \cdot M3]\left(\frac{K_1K_6 + K_4[S] + K_5[M1/2] + [S][M1/2]}{[M1/2][S]}\right)$$

$$[M1/2 \cdot S \cdot M3] = \frac{[M1/2][M3_{tot}][S]}{K_1K_6 + K_5[M1/2] + [S]([M1/2] + K_4)}$$

$$[M1/2 \cdot S \cdot M3] = \frac{\frac{[M1/2][M3_{tot}]}{K_4 + [M1/2]}[S]}{\frac{K_1K_6 + K_5[M1/2]}{K_4 + [M1/2]} + [S]}$$

$$rate = k_{chem}\left(\frac{\frac{[M1/2][M3_{tot}]}{K_4 + [M1/2]}[S]}{\frac{K_1K_6 + K_5[M1/2]}{K_4 + [M1/2]} + [S]}\right)$$

$$k_{cat,app} = \frac{k_{chem}[M1/2]}{K_4 + [M1/2]}$$ **Equation S5**

$$K_{M,app} = \frac{K_1K_6 + K_5[M1/2]}{K_4 + [M1/2]}$$ **Equation S6**

$$\left(\frac{k_{cat}}{K_M}\right)_{app} = \frac{k_{chem}[M1/2]}{K_1K_6 + K_5[M1/2]}$$ **Equation S7**

**Derivation of Michaelis–Menten kinetic equations according to Scheme 3**

$$rate = k_{chem}[M1/2 \cdot S \cdot M3^*]$$

$$[M3_{tot}] = [M3] + [S \cdot M3] + [M1/2 \cdot M3] + [M1/2 \cdot S \cdot M3^*]$$

$$K_1 = \frac{[M1/2][S]}{[M1/2 \cdot S]}\ ;\ [M1/2 \cdot S] = \frac{[M1/2][S]}{K_1}$$

$$K_2 = \frac{[M1/2][M3]}{[M1/2 \cdot M3]}\ ;\ [M1/2 \cdot M3] = \frac{[M1/2][M3]}{K_2}$$

$$K_3 = \frac{[M3][S]}{[S \cdot M3]}$$

$$K_4 = \frac{[M1/2][S\cdot M3]}{[M1/2\cdot S\cdot M3^*]}\,;\,[S\cdot M3] = \frac{K_4[M1/2\cdot S\cdot M3^*]}{[M12]}$$

$$K_5 = \frac{[M1/2\cdot M3][S]}{[M1/2\cdot S\cdot M3^*]}\,;\,[M1/2\cdot M3] = \frac{K_5[M1/2\cdot S\cdot M3^*]}{[S]}$$

$$K_6 = \frac{[M1/2\cdot S^*][M3]}{[M1/2\cdot S\cdot M3^*]}\,;\,[M3] = \frac{K_6[M1/2\cdot S\cdot M3^*]}{[M1/2\cdot S^*]}$$

$$K_7 = \frac{[M1/2\cdot S^*]}{[M1/2\cdot S]}$$

$$\frac{K_1K_6}{K_7} = K_3K_4 = K_2K_5$$

$$[M3_{tot}] = \frac{K_6[M1/2\cdot S\cdot M3^*]}{[M1/2\cdot S^*]} + \frac{K_4[M1/2\cdot S\cdot M3^*]}{[M1/2]} + \frac{K_5[M1/2\cdot S\cdot M3^*]}{[S]} + [M1/2\cdot S\cdot M3^*]$$

$$[M3_{tot}] = [M1/2\cdot S\cdot M3^*]\left(\frac{K_6}{K_7[M1/2\cdot S]} + \frac{K_4}{[M1/2]} + \frac{K_5}{[S]} + 1\right)$$

$$[M3_{tot}] = [M1/2\cdot S\cdot M3^*]\left(\frac{K_1K_6}{K_7[M1/2][S]} + \frac{K_4}{[M1/2]} + \frac{K_5}{[S]} + 1\right)$$

$$[M3_{tot}] = [M1/2\cdot S\cdot M3^*]\left(\frac{\frac{K_1K_6}{K_7}+K_4[S]+K_5[M1/2]+[S][M1/2]}{[M1/2][S]}\right)$$

$$[M12\cdot S\cdot M3^*] = \frac{[M1/2][M3_{tot}][S]}{\left(\frac{K_1K_6}{K_7}\right)+K_5[M1/2]+[S]([M1/2]+K_4)}$$

$$[M12\cdot S\cdot M3^*] = \left(\frac{[M12][M3_{tot}]}{K_4+[M12]}[S]\right)\Bigg/\left(\frac{\frac{K_1K_6}{K_7}+K_5[M12]}{K_4+[M12]} + [S]\right)$$

$$rate = k_{chem}\left(\frac{[M12][M3_{tot}]}{K_4+[M12]}[S]\right)\Bigg/\left(\frac{\frac{K_1K_6}{K_7}+K_5[M12]}{K_4+[M12]} + [S]\right)$$

$$k_{cat,app} = \frac{k_{chem}[M1/2]}{K_4+[M12]}$$

$$K_{M,app} = \frac{\frac{K_1K_6}{K_7}+K_5[M1/2]}{K_4+[M1/2]} = \frac{K_2K_5+K_5[M1/2]}{K_4+[M1/2]}$$

$$\left(\frac{k_{cat}}{K_M}\right)_{app} = \frac{k_{chem}[M1/2]}{K_2K_5+K_5[M1/2]}$$


1. Schuck, P. (1998) Sedimentation Analysis of Noninteracting and Self-Associating Solutes Using Numerical Solutions to the Lamm Equation. *Biophys. J.*, **75**, 1503-1512.
2. Schuck, P. (2000) Size-Distribution Analysis of Macromolecules by Sedimentation Velocity Ultracentrifugation and Lamm Equation Modeling. *Biophys. J.*, **78**, 1606-1619.
